# Comparing transmission reconstruction models with *Mycobacterium tuberculosis* whole genome sequence data

**DOI:** 10.1101/2022.01.07.475333

**Authors:** Benjamin Sobkowiak, Kamila Romanowski, Inna Sekirov, Jennifer L Gardy, James Johnston

**Author notes:** Department of Mathematics, Simon Fraser University, Burnaby, Canada (Benjamin Sobkowiak).

## Abstract

Pathogen genomic epidemiology is now routinely used worldwide to interrogate infectious disease dynamics. Multiple computational tools that reconstruct transmission networks by coupling genomic data with epidemiological modelling have been developed. The resulting inferences are often used to inform outbreak investigations, yet to date, the performance of these transmission reconstruction tools has not been compared specifically for tuberculosis, a disease process with complex epidemiology that includes variable latency periods and within-host heterogeneity. Here, we carried out a systematic comparison of seven publicly available transmission reconstruction tools, evaluating their accuracy in predicting transmission events in both simulated and real-world *Mycobacterium tuberculosis* outbreaks. No tool was able to fully resolve transmission networks, though both the single-tree and multi-tree input implementations of TransPhylo identified the most epidemiologically supported transmission events and the fewest false positive links. We observed a high degree of variability in the transmission networks inferred by each approach. Our findings may inform an end-user’s choice of tools in future tuberculosis transmission analyses and underscore the need for caution when interpreting transmission networks produced using probabilistic approaches.

## Introduction

Tuberculosis (TB), predominately caused by infection with *Mycobacterium tuberculosis* (Mtb), remains a major global public health concern, with an estimated 10 million people developing disease and 1.5 million deaths in 2020 ^1^. Accelerating TB burden reduction to meet EndTB’s elimination goals ^2^ will require broad improvements in our understanding of TB pathogenesis, along with improvements in access to point-of-care diagnostics, prevention, and effective treatment. It will also require a deeper understanding of Mtb transmission dynamics within different population settings – information critical to managing local outbreaks and developing data-driven prevention and control strategies based on an understanding of the clinical, social network, and environmental factors that drive onward transmission.

At the individual and population levels, identifying linked cases and broader transmission clusters has traditionally been done through a combination of DNA fingerprinting techniques and field-based epidemiological methods, such as contact tracing ^3^. While this approach can be effective in well-resourced settings with limited TB burden, it can be prohibitively labor-intensive in high transmission settings with limited laboratory and field epidemiology capacity. Moreover, contact tracing relies on the self-reporting of contacts ^4^ and is therefore inherently subjective and reliant on interviewee recall, which can be particularly problematic for an illness with a prolonged infectious period, such as TB; this can be limiting to the epidemiologists’ ability to accurately reconstruct full transmission histories.

Recently, Mtb whole genome sequencing (WGS) data has enabled finer-scale resolution of Mtb transmission events, although relating whole genome variation to direct transmission events can still be complex, particularly when clinical and epidemiological information, such as symptom onset date, bacillary load, and known contacts and places of social aggregation, are incomplete or unavailable. The most simplistic method is to set a threshold for the maximum level of genetic variation permitted to define transmission between samples, often using a single nucleotide polymorphism (SNP) threshold. This technique has been applied successfully in multiple settings to draw insights into transmission dynamics ^5,6^, but it often lacks sufficient resolution to determine the direction and timing of individual transmission events ^7,8^, and is further complicated by the lack of consensus on SNP thresholds ^5,9–11^.

A more sophisticated approach to transmission reconstruction using genomic data relies on phylogenetic trees. This method, phylodynamics ^12^, can characterize macro-scale network patterns ^13^ and estimate transmission events and timing for some pathogens, such as RNA viruses ^14^. However, in many bacterial pathogens, including Mtb, multiple factors complicate the use of phylodynamic approaches. These can include within-host evolution driven by variable latency periods or chronic infection, variable mutation rates, and mobile genetic elements ^14–16^. Clinical sampling can also affect our interpretation of genomic data - in cases where a host is infected with and capable of transmitting multiple lineages of Mtb, a sputum sample may contain or preferentially culture only a single lineage, leading to incorrect transmission inferences ^17,18^.

Recently, several groups have developed computational tools that combine genomic variation with an underlying epidemiological model (e.g. susceptible-infected-recovered (SIR) ^8,19^ or stochastic branching process ^16,20^ to estimate the probability of individual-level transmission events from genomic data alone. These tools take either a multi-sequence alignment or phylogenetic trees as input, along with other information, such as sampling date, and mainly employ a Bayesian Markov Chain Monte Carlo (MCMC) framework to account for the high dimensionality and computational complexity of the resulting models, as well as incorporating other epidemiologically derived parameters.

Many of these tools have been used in Mtb transmission studies ^21–23^; however, their performance on Mtb data has not yet been systematically compared. In this study, we evaluated seven model-based transmission inference approaches; seqTrack ^24^, TransPhylo single tree input ^16^, TransPhylo multiple tree input ^25^, Outbreaker2 ^26^, Phybreak ^20^, SCOTTI ^27^, and BEASTLIER ^19^. We first simulated ten Mtb outbreaks and the resulting genomic data and evaluated the accuracy of each method in identifying known *in silico* transmission links. Next, we ran each tool on real-world WGS data from clinical Mtb isolates collected in British Columbia, Canada, (BC) between 2005-2014. We compared the resulting high-probability transmission events inferred by each approach to known epidemiologically linked case-contact pairs and examined the methods’ different posterior estimates of key transmission parameters. Our results represent the first evaluation and comparison of multiple tools for reconstructing Mtb transmission networks using genomic data.

## Materials and methods

### Clinical sample information

Study data were collected in BC, a low TB burden region with a population of 5 million people and a TB incidence of 6 cases per 100,000 population ^28,29^. Mtb samples were obtained from the Public Health Laboratory (PHL) of the BC Centre for Disease Control (BCCDC), which maintains an archive of all Mtb culture-positive isolates from individuals diagnosed with TB in the province. From 2,915 culture-positive TB cases diagnosed between 2005-2014, genomic DNA was extracted from 2,290 isolates, one sequence per person, and analysed using MIRU-VNTR genotyping. Sample preparation, DNA extraction, and genotyping methods are described elsewhere ^30^. Ethics were obtained from the University of British Columbia (certificate H12-00910).

### Clinical sample sequence analysis

Whole genome sequencing was performed at the BC Genome Sciences Centre on the Illumina HiSeq platform on all isolates with a shared MIRU-VNTR genotyping pattern. This resulted in 1,014 high-quality whole genome Mtb sequences of 125bp paired-end reads, with an average depth of coverage of >100x. Sequencing reads were quality checked using the FastQC tool and processed to remove adapter sequences and low quality reads using Trimmomatic ^31^. Reads were mapped to the H37Rv reference strain (Genbank no.: NC_000962) using BWA **mem** ^32^.

Variant calling was conducted using the GATK suite of programs ^33^, including **HaplotypeCaller** and **GenetypeGVCFs**, with SNPs used in the subsequent analysis. Low confidence variants (phred quality score Q <20, read depth DP <5) and sites with a missing call (either through non-mapping of sequencing reads or low coverage) in >10% of isolates were removed. Heterozygous sites were called as the reference or alternative allele where ≥80% of mapped reads corresponded at these positions; otherwise, these sites were assigned as an ambiguous call ‘N’. Variants in repetitive regions, PE/PPE genes and at known resistance-conferring genes were also removed from the transmission analysis.

Putative transmission clusters were identified by grouping isolates with a shared MIRU-VNTR pattern, with preliminary analysis revealing that one cluster of 65 individuals, MCLUST012, was comprised of two distinct sub-clusters of 49 and 16 individuals and thus was split for further analysis. We tested transmission reconstruction tools on the resulting 12 clusters that we categorized as either small (four clusters between 5 and 9 isolates) or large (eight clusters with ≥ 10 isolates) and for which contract tracing data was present for at least one epidemiological linked pair of cases in the cluster.

### Phylogenetic methods

Timed phylogenies were produced for each MIRU-VNTR cluster separately with either BEAST v1.10.4 ^34^ (for BEASTLIER) or BEAST2 v2.6.3 ^35^ (for TransPhylo), calibrated at the tips by date of collection. A multiple sequence alignment of concatenated SNPs was used as input, with XML files modified manually to estimate the number of invariant sites. An appropriate substitution rate model was chosen for each cluster by producing a maximum likelihood tree using IQTREE ^36^ with 1000 bootstraps and applying the ‘ModelFinder’ algorithm ^37^ to find the model with the lowest Bayesian information criterion (BIC) score. We assessed the temporal signal in each cluster (the correlation between collection date and genomic distance) using TempEst ^38^. Where some temporal signal was found (*R*^*2*^ *>* 0.1), a strict molecular clock was used, otherwise we used a relaxed lognormal clock prior and supplied both models with an initial prior of 1×10^−7^ with a lognormal distribution, updated through MCMC iterations. Tree building parameters were further optimised for each cluster by conducting separate preliminary runs of 10^8^ MCMC iterations whilst varying the population model (constant, exponential, and Bayesian Skyline). Results were assessed using the posterior marginal likelihood estimates and run convergence in Tracer v1.7.1 ^39^, discarding the 10% of trees as the burn-in. The optimal tree building parameters chosen for each cluster are detailed in the supplementary materials along with the posterior estimates of cluster-specific substitution rate and tree height (**Supplementary Table S1**).

Final time-calibrated phylogenies were produced from BEAST2 runs of 10^9^ MCMC iterations with the optimised prior parameters, sampling every 10,000^th^ tree and discarding the first 10% of trees as the burn-in. Where a single phylogenetic tree was used as input for reconstruction models, a maximum clade consensus tree was obtained using Tree Annotator v2.6.3 ^35^ with median node heights. For analysis of multiple phylogenetic trees simultaneously with TransPhylo, a random selection of 50 trees was drawn from the posterior selection of 10,000 trees, discarding the 50% burn-in, using a custom bash script.

### Transmission network reconstruction models

We tested six tools for reconstructing transmission networks on the simulated outbreaks and real dataset of clinically derived Mtb isolates: seqTrack ^24^, TransPhylo ^16^, Outbreaker2 ^26^, Phybreak ^20^, SCOTTI ^27^, and BEASTLIER ^19^. There has been a recent extension to TransPhylo that allows for inference across multiple input trees sharing parameter estimates to account for uncertainty in the tree building process ^25^ - this was also tested and is referred to as TransPhyloMT. All approaches utilize genomic data, either directly as a multiple sequence alignment, or indirectly from a timed phylogenetic tree, as well as sampling dates. Input data types and epidemiological characteristics of the tested methods are shown in **Table 1**, along with specific model features, such as accounting for non-sampled hosts and within-host evolution.

**Table 1.**
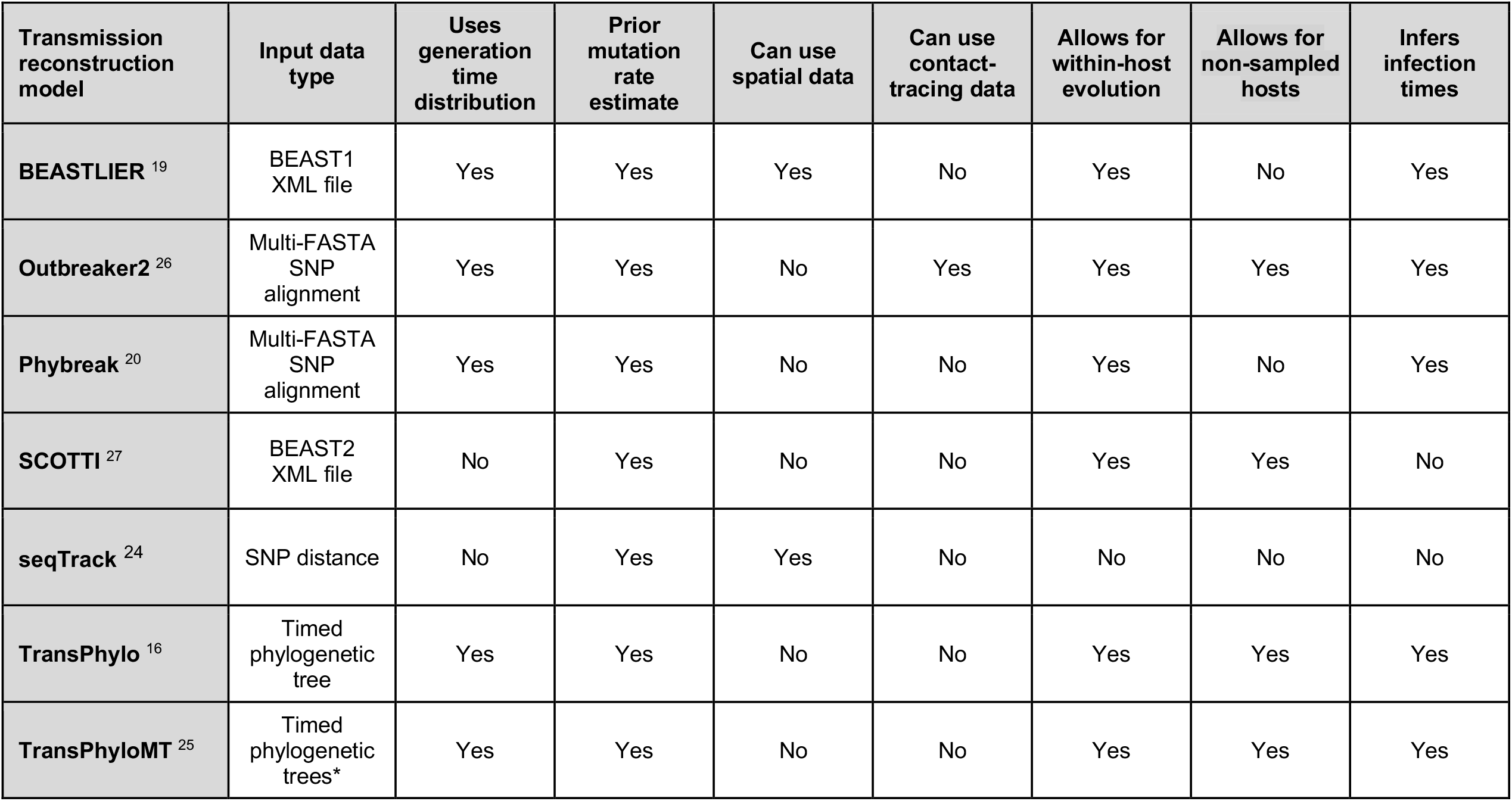
The transmission network reconstruction tools evaluated in this study, detailing the epidemiological features and input data type for each approach.

All tools allow specification of a prior estimate of either the generation time or host infectiousness interval to model the time interval between hosts becoming infected. For these measures, a long-tailed gamma distribution with parameters of shape = 1.3 and scale = 3.33 was chosen, previously used in transmission analysis of this TB population and in other settings with effective active case-finding strategies ^16,40^. Some methods also allow for a prior estimate of the sampling distribution - the time between a host becoming infected and the sample being taken; for these, we set a gamma distribution with shape and scale parameters of 1.1 and 2.5 ^16,40^. All methods, except BEASTLIER and seqTrack, allow for incomplete sampling by including unobserved hosts in predicted transmission networks (**Table 1**). Where permitted, we set a prior sampling proportion of around 80% complete sampling of TB cases in BC, a well-resourced setting with good active case finding ^41^. Full details for the model-specific inputs and prior parameter estimates can be found in the **Supplementary Materials**.

### Model performance on clinical data from British Columbia

In the absence of a gold-standard for confirmed transmission, the performance of each approach on clinical Mtb data was evaluated by calculating model sensitivity and positive predictive value (PPV) using the number of epidemiologically linked case-contact pairs that were correctly identified (**Supplementary Tables S2 and S3**). Case-contact pairs were defined as per ^41^ and included when both isolates are present in the same MIRU-VNTR defined cluster. Correct predictions (true positives) were classified as an inferred transmission event between a known case-contact pair identified with a probability of ≥ 0.5, with sensitivity defined as the proportion of the total case-contact pairs correctly identified for each cluster. Where multiple donor hosts were identified for a recipient strain with a probability of ≥ 0.5, only the highest probability link was considered. While not all true transmission events were captured in the case-contact pairs, meaning some true positive links will be misclassified as false positives, we attempted to account for models simply identifying a greater number of true transmission events through inferring a high number of links overall, including false positives, by calculating the PPV of each method. The PPV was calculated as the proportion of correct predictions in the total number of transmission links (P ≥ 0.5) inferred per cluster.

Additionally, we compared posterior estimates of transmission parameters to evaluate the credibility of the links predicted by each approach. Firstly, previous work has reported that a signal of direct transmission in Mtb is low divergence between sequences, with most transmission pairs differing by 0-5 SNPs ^5,7,42^. Therefore, while there may be some cases of sequences differing by more than 5 SNPs in direct recent transmission events, we can work under the assumption that most transmission will be between isolates with few SNP differences and an abundance of links between highly diverged sequences may indicate erroneous inferences. Five of the tested approaches, excluding seqTrack and SCOTTI, inferred an infection time for each host and thus we were able to calculate transmission intervals in these models, defined as the time between donor and recipient host infection. While this interval can be highly variable in TB, evidence suggests that most secondary transmission events occur within the two years after infection ^41,43^ and this should be reflected in the inferred transmission networks.

### Simulated Mtb transmission clusters

We simulated ten transmission clusters reflecting the epidemiology and sampling strategy in the clinical Mtb clusters used in this study using the ‘sim.phybreak’ function in Phybreak ^20^. Transmission clusters were simulated with an observed host number of 50, which we then sampled 6 years either side of the median sampling date to simulate an ongoing outbreak that was active before the date of the first observed host. To allow for the presence of non-sampled hosts within clusters, we randomly down-sampled the observed hosts within each cluster by 20% to simulate incomplete sampling, which resulted in a final sample number in clusters of between 28 and 40 observed hosts.

Epidemiological parameters for the simulated clusters were chosen to reflect clinical Mtb clusters, thus we chose the same gamma generation and sampling time distributions as described previously. The sequence length was set as length of the H37Rv Mtb reference stain at 4.4M base pairs and we allowed for within-host evolution at the rate of 1.48 as previously estimated ^16,40,44^ and a mutation rate of 0.5 SNPs per genome per year, taken from estimates in our clinical data using BEAST2 v2.6.3 ^35^, which is in line with previous estimates of the Mtb mutation rate ^45^. The simulator takes these epidemiological estimates and produces simulated sequence data and transmission networks, including the time and direction of transmission.

Simulated genome sequences and sampling dates of the observed hosts were used as input for each tested approach, with the resulting predicted transmission events compared against known transmission to evaluate model sensitivity (the proportion of the true transmission links correctly identified) and PPV (the correct transmission links as a proportion of the total inferred transmission links (true positives + false positives)). Additionally, simulated transmission networks included information on who-infected-whom and so sensitivity and PPV for inferring the correct direction of transmission was also calculated for each model.

## Results

### Simulated transmission clusters

We first assessed the performance of the described methods by identifying the number of transmission events correctly predicted in ten simulated Mtb outbreaks. **Figure 1A** shows the sensitivity and PPV of each approach for all simulated clusters for transmission links with a probability of ≥ 0.5, irrespective of the direction of transmission (i.e., if host i transmitted to host j in the true transmission network, a prediction of i -> j or j -> i would be scored as correct). Outbreaker2 correctly predicted the highest proportion of the true transmission events with a median sensitivity of 0.5 (IQR 0.37 – 0.58), followed by TransPhylo (sensitivity 0.4, IQR 0.35 – 0.44), though Phybreak and TransPhyloMT achieved comparable scores (median 0.4, IQR 0.36 – 0.49 and median 0.38, IQR 0.35 – 0.43 respectively). BEASTLIER, seqTrack and SCOTTI performed poorly in predicting transmission in the simulated clusters and, notably, SCOTTI did not infer any transmission links with a probability of ≥ 0.5 in any cluster (Supplementary data, Table S2). There were marked differences in the PPV of each approach when predicting correct transmission links in simulated clusters, with TransPhyloMT achieving the highest PPV (median 0.68, IQR 0.62 – 0.72), followed by TransPhylo (median 0.58, IQR 0.48 – 0.66). The high PPV of TransPhyloMT compared to the other higher sensitivity approaches was due to fewer false positive transmission links predicted with a probability of ≥ 0.5 (Supplementary data, Table S2).

**Figure 1.**
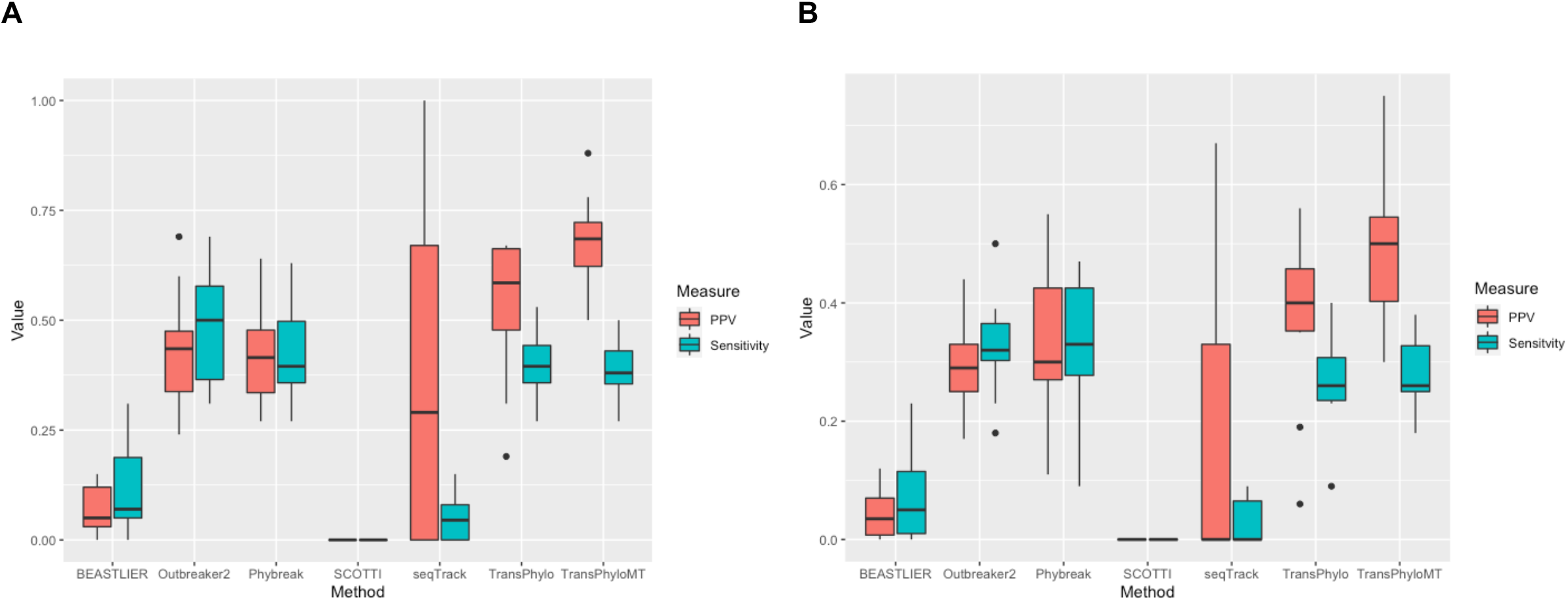
Boxplots showing the sensitivity (blue boxes) and positive predictive value (PPV) (red boxes) of each transmission reconstruction model for predicting known transmission events with a probability of ≥ 0.5 in ten simulated TB clusters.

All models predicted fewer correct transmission links when the direction of transmission was considered (i.e., i -> j ≠ j -> i) (**Figure 1B)**. Phybreak achieved the highest sensitivity, predicting around a third of true directional transmission links (median 0.33, IQR 0.27 – 0.42), followed by Outbreaker2 (median 0.32, IQR 0.3 – 0.36) and both TransPhylo and TransPhyloMT (median 0.26, IQR 0.24 – 0.31, and median 0.26, IQR 0.25 – 0.33). The PPV of TransPhyloMT was again highest when identifying the direction of transmission, with typically around half of the inferred transmission events predicted using this approach being a confirmed link (median 0.5, IQR 0.41 – 0.54).

### Clinical M. tuberculosis data from British Columbia

We next applied each method to real-world WGS data. We observed a high degree of variation in transmission networks inferred by each method, reflecting the different underlying model algorithms and parameters employed by each approach. An example of the differences in consensus transmission networks is shown in **Figure 2**, which illustrates transmission events inferred by two methods, Phybreak and TransPhylo, for cluster MCLUST004 (N = 9 cases).

**Figure 2.**
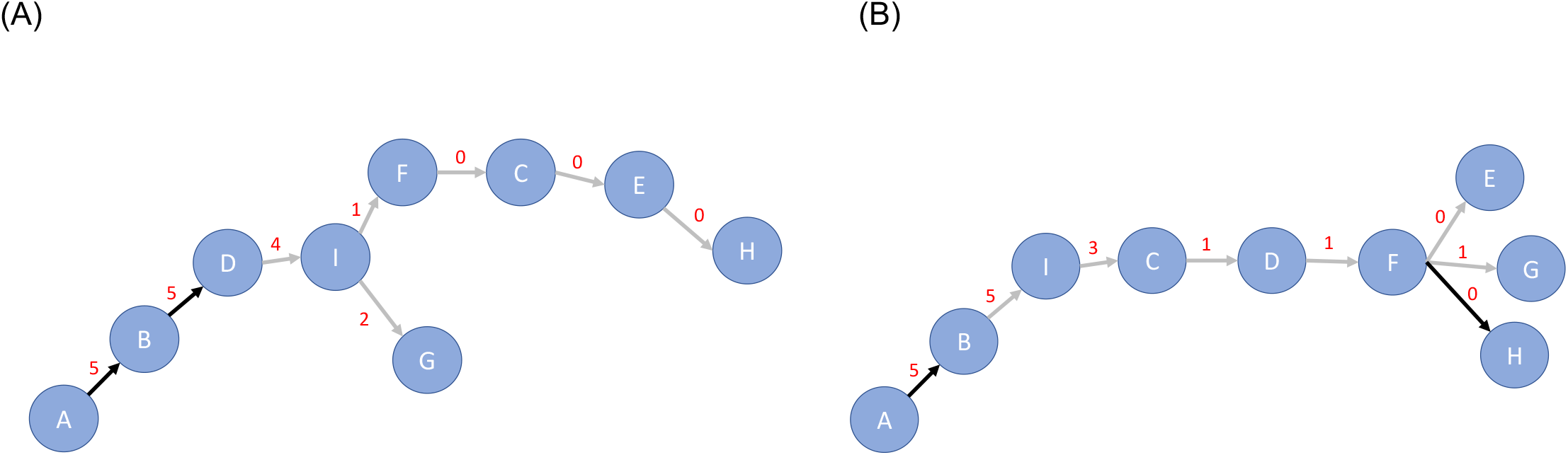
Example transmission networks inferred using two of the test methods, (A) Phybreak and (B) TransPhylo single tree, for the clinical BC transmission cluster, MCLUST004. Blue nodes represent the sampled hosts with sample names illustrated by letters A - I, black edges represent links with a posterior probability of ≥ 0.5, and grey edges represent links < 0.5, with the SNP distance between sampled hosts in red on edges.

**Table 2** shows the PPV and sensitivity of each model for predicting transmission within the clinical Mtb clusters, as both a total for all clusters and separated by small (N < 10) and large (N ≥ 10) clusters. Overall, the highest number of the epidemiologically linked case-contact pairs were identified with TransPhylo (20/120; sensitivity 17%) followed by TransPhyloMT (18/120; sensitivity 15%). SCOTTI again performed poorly, predicting just a single case-contact pair in the resulting transmission links (sensitivity 1%). There were clear differences between model sensitivity in the large and small clusters, with the best scoring model, Phybreak, identifying just 16/106 (15%) case-contact pairs in large clusters, while TransPhyloMT correctly identified 6/14 (43%) case-contact pairs in the small clusters. PPV was also highest using TransPhyloMT for all clusters, as well as when considering small and large clusters separately, with 18/63 (29%) epidemiologically linked case-contact pairs inferred as high-confidence transmission links, suggesting this method will infer the lowest number of false positive transmission events.

**Table 2.** The positive predictive value (PPV) and sensitivity of transmission pairs and epidemiologically linked pairs identified in clinical Mtb isolates from British Columbia.

| Transmission reconstruction model | All clusters |  | Small Clusters (N < 10) |  | Large Clusters (N ≥ 10) |  |
| --- | --- | --- | --- | --- | --- | --- |
|  | PPV | Sensitivity | PPV | Sensitivity | PPV | Sensitivity |
| **BEASTLIER** <sup>19</sup> | 15/320 (5%) | 15/120 (13%) | 5/25 (20%) | 5/14 (38%) | 10/295 (3%) | 10/106 (9%) |
| **Outbreaker2** <sup>26</sup> | 14/133 (11%) | 14/120 (12%) | 1/10 (10%) | 1/14 (7%) | 13/123 (11%) | 13/106 (12%) |
| **Phybreak** <sup>20</sup> | 17/143 (12%) | 17/120 (14%) | 1/9 (11%) | 1/14 (7%) | 16/134 (12%) | **16/106 (15%)** |
| **SCOTTI** <sup>27</sup> | 1/4 (25%) | 1/120 (1%) | 0/0 (0%) | 0/14 (0%) | 1/4 (25%) | 1/106 (1%) |
| **seqTrack** <sup>24</sup> | 8/80 (10%) | 8/120 (7%) | 1/4 (25%) | 1/14 (7%) | 7/76 (9%) | 7/106 (7%) |
| **TransPhylo** <sup>16</sup> | 20/135 (15%) | **20/120 (17%)** | 5/14 (36%) | 5/14 (38%) | 15/119 (13%) | 15/106 (14%) |
| **TransPhyloMT** <sup>25</sup> | **18/63 (29%)** | 18/120 (15%) | **6/11 (55%)** | **6/14 (43%)** | **12/52 (23%)** | 12/106 (11%) |

Closer inspection revealed that all models predicted a low proportion of the case-contact pairs in the largest two clusters in the population, MCLUST001 and MCLUST002 (**Supplementary Table S3**), this though may be due to the complicated epidemiology in these two large outbreaks ^46^ and so we can be less confident of the validity of the case-contact pairs. Indeed, removing these two clusters from the analysis significantly increased the sensitivity scores of most models, with Phybreak now correctly predicting 13/33 (39%) of case-contact pairs in the remaining large clusters.

Posterior estimates of SNP distance and transmission interval between hosts in predicted direct transmission events revealed differences between tested approaches. The median SNP distance in high probability transmission links between observed hosts was low for all models, ranging from 0 SNPs (IQR 0 – 0 SNPs) in seqTrack, which only predicted transmission events between identical sequences, to 5 SNPs (IQR 2 – 9 SNPs) for BEASTLIER (**Figure 3A**). BEASTLIER and Phybreak also predicted several transmission events between relatively divergent isolates (> 30 SNP differences), which may be due to these models not allowing for non-sampled hosts, resulting in incorrect direct links between hosts when there are many unobserved cases in the real transmission chain. **Figure 3B** illustrates the differences in transmission interval estimated between observed hosts in direct transmission events for models that inferred infection times. We can see that all tested approaches estimate that the majority of secondary cases are within the first two years after donor host infection (BEASTLIER median 0.2 years, IQR 0.2 – 0.5, to Phybreak median 1.1 years, IQR 0.5 – 2.1). Phybreak and BEASTLIER predict some high-probability transmission events between hosts where the infection time is many years, with a maximum value of 8.8 years between host infection times using BEASTLIER. While long periods between transmission events have been observed in Mtb due to latency of disease onset, the links inferred here are not supported by epidemiological evidence and as these only occur in models only allowing transmission between observed hosts, these are likely an artifact of the model assumptions.

**Figure 3.**
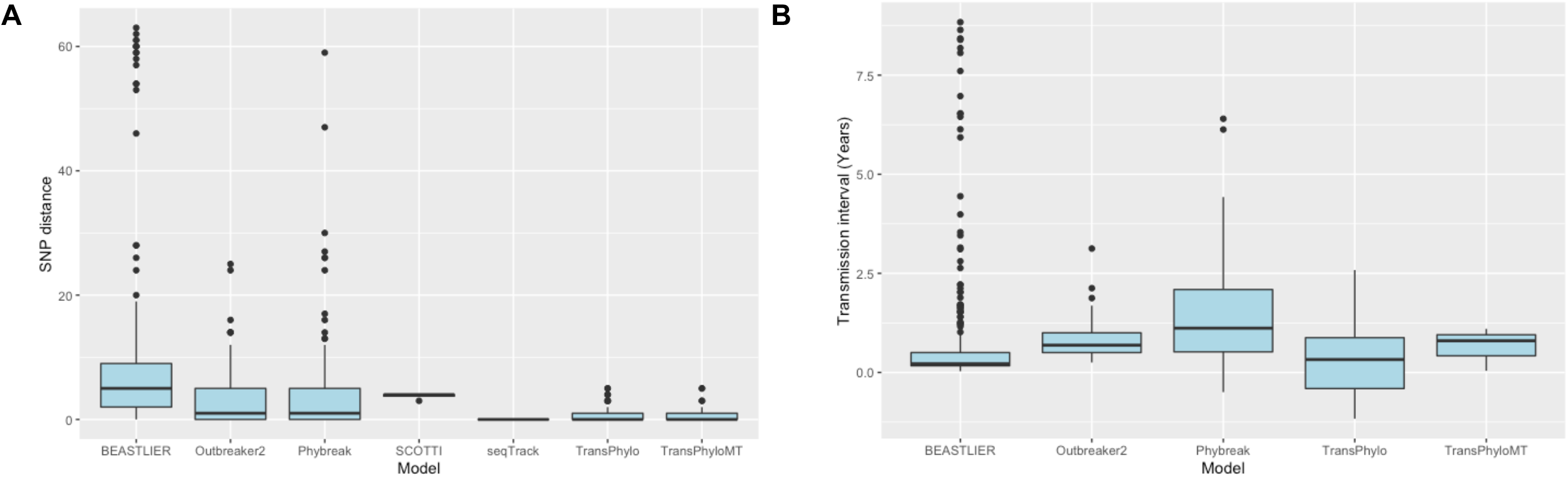
Boxplots of the transmission parameters estimated by each tested method from high probability (*P ≥* 0.5) transmission events between sampled clinical Mtb isolates from BC. (A) the SNP distance between observed hosts, and (B) the transmission interval between infection times of observed hosts.

### Sensitivity analysis

To assess the impact of the setting the accepted probability threshold of transmission links at P ≥ 0.5 in the main analysis, we re-calculated the sensitivity and PPV of each approach for the clinical BC MIRU-VNTR clusters at a higher and lower probability threshold of P ≥ 0.75 and P ≥ 0.25. As expected, we found an increase in the sensitivity of most methods when decreasing the probability threshold to ≥ 0.25, and a decrease in sensitivity at P ≥ 0.75 (**Supplementary figure S1**). These differences were most pronounced in Phybreak, TransPhylo, and TransPhyloMT, where a lower probability threshold allowed for a greater number of case-contact pairs to be identified. Lowering the probability threshold also increased the overall number of accepted transmission links with each tested model, which may also increase the number of false positive calls. The PPV of both TransPhylo and TransPhyloMT decreased markedly at a probability threshold of ≥ 0.25 compared to ≥ 0.5, with a greater number of false positive transmission links identified (**Supplementary figure S2**). The PPV of seqTrack and Outbreaker2 increased at P ≥ 0.25 compared to P ≥ 0.5, though the relative difference between these values was small. Interestingly, we saw a decrease in PPV for all tools at the ≥ 0.75 probability threshold compared to ≥ 0.5 as the reduction in the number true positive calls was greater than the decrease in false positives at this higher probability threshold.

## Discussion

We present the first systematic comparison of the available methods for reconstructing TB transmission networks using whole genome sequence data, using input data from both simulated outbreaks and real-world Mtb transmission clusters. Our results demonstrate that the choice of analytical tool can impact the structure and accuracy of predicted transmission events. While it is difficult to infer full Mtb transmission networks using only genomic and temporal data owing to the low diversity characteristic of TB outbreaks, we have shown certain models performed markedly better than others in both real-world and simulated data. As such, the results presented here can help to guide the choice of methodology for Mtb transmission investigations.

In simulated outbreaks, Outbreaker2 predicted the highest number of ‘true’ transmission events, with an average of half the links between hosts in each cluster correctly identified, and a third predicted with the correct direction of transmission. Phybreak, TransPhylo and TransPhyloMT achieved comparable results for identifying known transmission between hosts, though the overall number of high probability links inferred by Outbreaker2 and Phybreak was higher than with TransPhylo and TransPhyloMT. This greatly increased the number of false positive transmission events predicted by both Outbreaker2 and Phybreak, which resulted in the PPV of these tools being significantly lower than TransPhylo and TransPhyloMT. This is also seen in the specificity of each approach on the simulated clusters; while the differences are not as large, TransPhylo and TransPhyloMT both outperformed Phybreak and Outbreaker2 (**Supplementary Figure S3**). Therefore, these results indicate that TransPhylo, particularly TransPhyloMT, may be the most effective tool for identifying transmission events between sampled hosts in a population, while reducing the number of spurious links between hosts that are wrongly predicted with a high probability.

A limitation in this study was an absence of a gold-standard for transmission within the clinical TB population. We were able to provide strong evidence of ‘correct’ links by including several epidemiologically linked case-contact pairs in our analysis, but these data were drawn from contact tracing questionnaires where participants may not produce a complete network of potential sources of infection. Nonetheless, we hypothesized that the presence of epidemiological and genomic linkage between hosts increased the likelihood of these links being true transmission events and thus these links were used as a measure of ‘ground-truth’ in analysis of the clinical TB data.

Unfortunately, the ability of all tested models to identify case-contact pairs was relatively low, with TransPhylo performing best but only finding 17% of epidemiologically supported links. The poor performance of models appears to be driven by the low number of case-contact pairs inferred in the largest two clinical transmission clusters, MCLUST001 and MCLUST002. These clusters were part of two separate TB outbreaks with complex demography, that included people experiencing homelessness and poly-substance use ^47,48^. The disparity between the inferred transmission networks and the reported case-contact information may be due to difficulty in contact tracing in these settings, decreasing the likelihood of the contact data containing the true transmission links. Indeed, after removing MCLUST001 and MCLUST002 from analysis, the overall performance of most tools increased, with Phybreak identifying 39% and TransPhylo 33% of the remaining epidemiologically linked case-contact pairs (**Supplementary Table S3**).

In this study, TransPhyloMT achieved the highest PPV in clinical data, with epidemiologically linked case-contact pairs constituting 29% of the total high probability transmission events inferred, rising to 55% in small clusters. In contrast, we found the number of transmission events inferred by SCOTTI in the clinical Mtb data to be particularly low, with only four direct transmission links inferred across all 12 clusters, and only a single epidemiologically confirmed case-contact pair detected. The reason for this may be two-fold; SCOTTI was not primarily developed to model transmission in Mtb populations that are often characterised by few clonal lineages and low genomic variation. In addition, we used a probability threshold for an accepted transmission link of ≥ 0.5, while the majority of the reported links for real-life outbreaks tested in the original publication of the model were below this value ^27^. Indeed, when we decreased the threshold value to ≥ 0.25, we observed a vast increase in both the overall number of predicted transmission events between sampled hosts (54 at ≥ 0.25 compared to 4 at *P ≥* 0.5), as well as an increase in the number of identified epidemiologically linked pairs (7/120 at ≥ 0.25 compared to 1/120 at *P ≥* 0.5), (**Supplementary table S4**). However, even at this lower threshold, there were fewer epidemiologically linked pairs predicted with SCOTTI when compared to other models at higher accepted probability threshold.

No single transmission network was fully resolved with any tested model; Phybreak identified the highest proportion of known transmission events in a single cluster, while TransPhyloMT achieved the highest PPV within a cluster. To increase the number of correctly inferred transmission events, the threshold to accept an inferred link between hosts could be reduced, but this will lead to a greater number of false positive links. Previous studies have used these inference tools to gain insights in the Mtb transmission dynamics notably identifying individual transmission events and increased transmissibility within populations (e.g., ^21,23,25^), by reconstructing transmission histories based on accepting or rejecting links at a set probability threshold. We have demonstrated that there can be doubt in the inferred transmission links, even with the best performing models, and as such it may be valuable to provide the probability of linkage between hosts, rather than a binary classifier probability threshold for accepting transmission.

As computational models for transmission inference are being improved to better reflect the characteristics of disease outbreaks and incorporate more realistic epidemiological parameters, we suggest that a TB-focused methodology will likely deliver more accurate transmission networks than tools calibrated to be applicable across a range of pathogen genomes. This is reflected in the results shown here, where the best performing models, Phybreak, Outbreaker2, and TransPhylo, have been developed to account for the complex epidemiology of infectious diseases like tuberculosis, including handling within-host evolution and non-complete outbreak sampling ^16,20,26^. Furthermore, some modifications could be implemented to existing tools to improve the accuracy of inferred transmission networks. For example, other forms of genomic variation could be used to increase the observed variation between strains, such as small insertions and deletions (INDELs) that are often found in single isolates ^49^. This may improve resolution in instances when multiple isolates are separated by very few SNPs, typical in highly clonal Mtb populations. Additionally, Outbreaker2 can be supplied with custom prior likelihood and movement functions to adapt the underlying model, as well as accepting contact network data, and these modifications can be expected to improve the accuracy of output networks when appropriately specified *a priori*.

Together with the performance of each model for predicting transmission events, a further consideration when running these tools is the processing complexity and runtime, which may influence the decision on the model to employ. This is particularly true if the aim is to use these approaches for real-time surveillance of Mtb transmission or in settings with limited computational resources. Whilst all the tested tools ran successfully using a desktop computer with a 2.6GHz processor and 16GB memory, the runtime of each approach varied considerably (**Supplementary Table S5**). The fastest tool tested was seqTrack, with networks computed almost instantaneously for the largest clinical Mtb cluster MCLUST002 (N = 74 cases). This is due to the simplicity of this approach compared to other models, which employ a Bayesian framework that must run over millions of MCMC iterations for chain convergence. However, this method may not provide accurate transmission networks in populations with within-host evolution such as Mtb and incomplete sampling, as shown by our results. While we observed reasonable inference of transmission using both TransPhylo approaches, these models took considerably longer to complete in cluster MCLUST002 (∼19 hours for TransPhylo MT, including phylogeny construction) and this would need to be taken into consideration when selecting this tool. These results only show limited comparisons of the computational resources required by each approach and more work would be required to test the impact of running fewer iterations on accuracy with different computational infrastructure to properly benchmark each approach.

While this study is not an exhaustive analysis of all transmission network reconstruction methodologies (e.g. ^50^), we have presented results from selected models that have been used for investigating pathogen transmission histories and where a software package was freely available for implementation. In addition to the methods presented here, we also attempted to reconstruct transmission using BadTrIP ^51^, which also employs Bayesian inference to infer transmission networks from sequence data and accounts for site polymorphisms, transmission bottlenecks and sequencing error. This model, though, failed to reach convergence in five of the eight large clinical clusters/sub-clusters of BC isolates when allowing to run for 5×10^8^ MCMC iterations, and so was not included in this study.

In conclusion, we have systematically compared seven tools for reconstructing transmission using genomic data to assess their utility for TB transmission analysis. The results presented here indicate that the transmission networks produced by each method can disagree markedly due to the differences in the underlying models. While there are limitations in the accuracy of the transmission histories provided by all models, we found that the multi-tree implementation of TransPhylo achieved the most effective balance of identifying a relatively high number of true transmission links while predicting fewer false positive events in both simulated and real-life clinical Mtb clusters. This approach could be applied to gain some insights into Mtb transmission dynamics using sequence data in settings with limited contact network information. These findings can improve investigations into Mtb outbreaks and transmission dynamics and highlight further areas of research to advance methods for transmission network reconstruction.

