## Supplementary material for "Comparing transmission reconstruction models with *Mycobacterium tuberculosis* whole genome sequence data": Supplementary Materials.docx

‡. Present address. Department of Mathematics, Simon Fraser University, Burnaby, Canada

**Supplementary Methods**

The following are model specific methods and parameter choices for each of the tested models used to reconstruct transmission in simulated and clinical clusters of *Mycobacterium tuberculosis*.

*BEASTLIER*

BEASTLIER ^1^ works by modifying the input XML file used by BEAST v1.8.4 or later to estimate transmission probabilities between hosts to produce a consensus transmission network, along with the phylogenetic tree output from BEAST. The resulting transmission network will include the highest probability links between all hosts and does not allow for non-sampled hosts in the inferred network.

A python2 script generates a new input file for BEAST and requires three input files: the standard XML file produced by BEAUTi, a CSV file containing the name and date of becoming non-infectious for each host, and a CSV file containing the sequence name, host name and collection dates of each sequence. The date that a host became non-infectious was set as three months past the collection date (using collection date as a proxy for the date of treatment onset) ^2^, and as we had only one sequence per host, a dummy host name was supplied to avoid replication of sequence names. The input XML file was produced with BEAUTi as described in the main manuscript.

The period of infectiousness, defined as the time between a host becoming infectious and end of infectiousness, was given a gamma distribution with shape and scale parameters of 1.1 and 2.5, reflecting the BCCDC guidelines for contact investigation stating that an infectious period of at least three months prior to diagnosis is projected, though with a reasonably high active case finding in our population suggesting few cases will stay infectious for a prolonged period ^3^. The period of latency, defined as the period between a host being infected and becoming infectious, was also given a gamma distribution with shape and scale parameters of 1.3 and 3.33, allowing for a variable latency period with a long-tailed distribution but with the incubation period of most cases falling with the first 6 months to two years, as is estimated in low-burden settings ^4–6^. The within-host demographic model was kept retained from the model described in Hall *et al* ^1^. MCMC chains were run for 100 million iterations or until model convergence (ESS score > 200), logging every 10,000 iterations.

*Outbreaker2*

Outbreaker2 ^7^ is an update of the Outbreaker tool ^8^ implemented in R, which uses a Bayesian framework for reconstructing outbreaks for sequence data but now allows for modulization with user-defined models for the likelihood, priors and movement estimation. While the option to customize these components was available, we ran this analysis with the pre-defined models in Outbreaker2. The updated model allows for within-host evolution and non-sampled hosts. We supplied the tool with sequence data, as a multi-sequence alignment in FASTA format, and collection dates. Prior epidemiological parameters for the generation time distribution, colonization (sampling) time distribution, sampling proportion and mutation rate can be specified with a fixed value or updated through MCMC runs. For our analysis, the generation time distribution (gamma distribution with shape = 1.3 and rate = 0.3), sampling time distribution (gamma distribution with shape = 1.1 and rate = 0.4), and initial mutation rate estimated from each cluster’s phylogeny in BEAST (**Table S1**), updated through MCMC iterations ^9,10^. We estimated up to 80% of cases would be captured in our dataset and thus we set a sampling proportion with informative beta priors of 8 and 2, updated through MCMC runs. The model was run for 10^5^ MCMC chains, sampled every 10^3^ iterations.

*Phybreak*

Phybreak ^11^ is implemented in R that uses a Bayesian inference of both transmission and phylogenetic trees simultaneously. The tool uses sequence data, as a multi-sequence FASTA file, and collection date to produce a ‘phybreakdata’ object as input to the main algorithm, where you can supply user-defined prior parameters. We again set the generation time distribution (gamma distribution with shape = 1.3 and rate = 0.3), sampling time distribution (gamma distribution with shape = 1.1 and rate = 0.4), and initial mutation rate estimated from each cluster’s phylogeny in BEAST (**Table S1**), updated through MCMC iterations ^9,10^. We set a within-host effective population size with a slope rate of 1.48 x t (time after infection), based on previous estimates ^9^. The model was run for 10^5^ MCMC chains, sampled every 10^3^ iterations.

*SCOTTI*

SCOTTI ^12^ is available as an extension package in BEAST2 and estimates transmission probabilities between samples, along with sampling from a pool of generic non-sampled hosts (with the number specified by the user), to reconstruct transmission trees using a structed coalescent model ^12^. A python script is used to create a modified XML file that can be run with BEAST2 and requires four input files: a FASTA sequence file, a CSV file of the sampling dates, a CSV file of the host names, and a CSV of infection intervals for each host. Again, in our dataset there was one sequence per host so dummy host names were used. The infection interval spans the earliest time a host is infectible to the latest time in which they are infectious. We allowed for periods of latency where an individual may be infected before the disease becomes active by setting the earliest time for each host to be able to be a recipient of transmission as three years prior to the earliest sample collection date, and the end date of infectiousness to be three months after the collection date ^2^.

We allowed SCOTTI to include non-sampled hosts in the resulting transmission trees by providing the maximum number of hosts (observed and unobserved) in the tree. Again, as we estimate that we have captured a high proportion of the cases in our population (70-80% of all cases), we set the maximum number of hosts as 1.5x the number of observed hosts. We also set the mutation model to be employed for each cluster (Table S1) and an estimation of the number of invariable sites per nucleotide. MCMC chains were run for 100 million iterations to allow for model convergence, logging every 10,000 iterations.

*SeqTrack*

SeqTrack ^13^ uses a graph approach to reconstruct transmission trees directly from sequence data, considering ancestry between all sampled hosts, not allowing for non-sampled individuals in the network. Where there is equal likelihood of transmission between multiple hosts, these conflicts are resolved by finding the most parsimonious link when likelihood of the genetic differentiation using collection dates. This tool is run in R and uses sequence data, as a multi-sequence FASTA file, and collection dates. We also supplied a mutation rate (estimated from each cluster’s phylogeny in BEAST (**Table S1**)), and sequence length (4,411,532bp).

*TransPhylo*

TransPhylo ^9^ is a package in R that employs a stochastic branching process in a Bayesian framework for reconstructing transmission networks via a Monte Carlo Markov chain from genomic data, notably allowing for within-host evolution and incomplete sampling of the population by including non-sampled hosts into resulting networks. This approach requires a timed phylogenetic tree as input, with sampled hosts corresponding to the tips. Prior epidemiological parameters for the generation time distribution, sampling time distribution, sampling proportion and within-host effective coalescence rate can be specified and either given a fixed value or be updated through MCMC runs. For our analysis, the generation time distribution (gamma distribution with shape = 1.3 and rate = 0.3), sampling time distribution (gamma distribution with shape = 1.1 and rate = 0.4), and a fixed within-host effective coalescence rate (100/365) were chosen based on a previous analysis of the a transmission cluster in this study population ^9^. We again estimated up to 80% of cases would be captured in our dataset and thus we set a sampling proportion with informative beta priors of 8 and 2, updated through MCMC runs. The algorithm was run for 10^5^ MCMC chains, with the end date of the clusters set as three years past the latest sample collection date in the cluster to allow for some uncaptured transmission from later samples to non-sampled hosts to be potentially resolved.

Additionally, there is an extension to the original TransPhylo package that aims to improve the inferred transmission network by accounting for some uncertainty in the input phylogenetic tree by instead running simultaneous inferences with the input trees drawn from a random distribution of posterior trees produced by BEAST tools ^14^. Different model parameters can be shared across runs, which allows for better mixing of the MCMC iterations. To run this method, which we refer to as TransPhyloMT, we selected 50 input phylogenetic trees drawn randomly from the posterior selection of inferred phylogenies (after a 10% burn-in) produced by BEAST2 for each cluster using a custom bash script. All model parameters were kept the same as the single tree implementation, with parameter sharing, and transmission probabilities calculated as the mean probability of host-host transmission across the 50 runs.
