## Supplementary material for "Comparing transmission reconstruction models with *Mycobacterium tuberculosis* whole genome sequence data": Table S1.pdf

**Table S1** – Tree building parameters used to produce time-calibrated phylogenies using both **BEAST1** and **BEAST2**. The optimal substitution model was found using the 'ModelFinder' test in **IQTREE** and the root-tip correlation calculated in **TempEst** using untimed maximum likelihood phylogenies produced with **IQTREE** and collection dates. Substitution rates are posterior estimates from the posterior **BEAST2** trees, calculated in **Tracer**.

| Cluster Name | Substitution model | Root to tip correlation | No. of variants | Population model | Clock rate | Estimated substitution Rate |
|---|---|---|---|---|---|---|
| **MCLUST001** | HKY | 0.189 | 2250 | Constant | Strict | 1.23E-07 |
| **MCLUST002** | HKY | 0.301 | 2274 | Constant | Strict | 1.01E-07 |
| **MCLUST003** | HKY | 0.422 | 548 | Constant | Strict | 1.55E-07 |
| **MCLUST004** | HKY | 0.226 | 825 | Constant | Strict | 4.21E-07 |
| **MCLUST006** | HKY | 0.120 | 904 | Constant | Strict | 9.98E-07 |
| **MCLUST007** | HKY | 0.316 | 614 | Constant | Strict | 1.18E-06 |
| **MCLUST008** | HKY | 0.417 | 1306 | EBSP | Strict | 1.44E-07 |
| **MCLUST012_1** | HKY | 0.196 | 1244 | Constant | Strict | 1.17E-07 |
| **MCLUST012_2** | HKY | 0.119 | 1244 | Constant | Strict | 1.58E-07 |
| **MCLUST052** | HKY | 0.541 | 847 | Constant | Strict | 1.83E-07 |
| **MCLUST058** | HKY | 0.102 | 1257 | Constant | Strict | 4.12E-07 |
| **MCLUST134** | HKY | 0.338 | 935 | Constant | Strict | 1.73E-07 |
