## Supplementary material for "Comparing transmission reconstruction models with *Mycobacterium tuberculosis* whole genome sequence data": Table S4.pdf

**Table S4** – Model performance of SCOTTI using a lower probability threshold (≥ 0.25) for accepting transmission links in inferred transmission networks for clinical MIRU-VNTR *Mycobacterium tuberculosis* clusters. Sensitivity and positive predictive value (PPV) are calculated as detailed in the main text.

| Cluster Name | No. epi-associated links between sampled hosts | No. Inferred links | No. correctly predicted links | PPV | Sensitivity |
|---|---|---|---|---|---|
| **MCLUST001** | 58 | 12 | 3 | 25% | 5% |
| **MCLUST002** | 15 | 14 | 0 | 0% | 0% |
| **MCLUST003** | 16 | 3 | 1 | 33% | 6% |
| **MCLUST004** | 4 | 1 | 0 | 0% | 0% |
| **MCLUST006** | 5 | 0 | 0 | 0% | 0% |
| **MCLUST007** | 4 | 2 | 1 | 50% | 25% |
| **MCLUST008** | 7 | 8 | 1 | 13% | 14% |
| **MCLUST012_1** | 5 | 7 | 2 | 29% | 40% |
| **MCLUST012_2** | 1 | 1 | 0 | 0% | 0% |
| **MCLUST052** | 3 | 2 | 0 | 0% | 0% |
| **MCLUST058** | 1 | 2 | 1 | 50% | 100% |
| **MCLUST134** | 1 | 2 | 0 | 0% | 0% |
| **Total** | **120** | **54** | **9** | 17% | 8% |
