## Supplementary material for "Comparing transmission reconstruction models with *Mycobacterium tuberculosis* whole genome sequence data": Table S5.pdf

**<u>Table S5</u>** – Run times of tested methods on clinical MIRU-VNTR *Mycobacterium tuberculosis* cluster MCLUST003, comprising 38 host, using a desktop computer with a 2.6GHz processor and 16GB memory. The number of iterations for Bayesian approaches using MCMC methods is specified in the Supplementary Methods.

| **Method** | **Run time (hours)** |
|---|---|
| BEASTLIER | 49 |
| Outbreaker2 | 1.4 |
| Phybreak | 2 |
| SCOTTI | 65 |
| seqTrack | < 1 |
| TransPhylo | 1.5 |
| TransPhyloMT | 32 |
