## Supplementary figures and images for "Comparing transmission reconstruction models with *Mycobacterium tuberculosis* whole genome sequence data"

### Figure S1.tiff

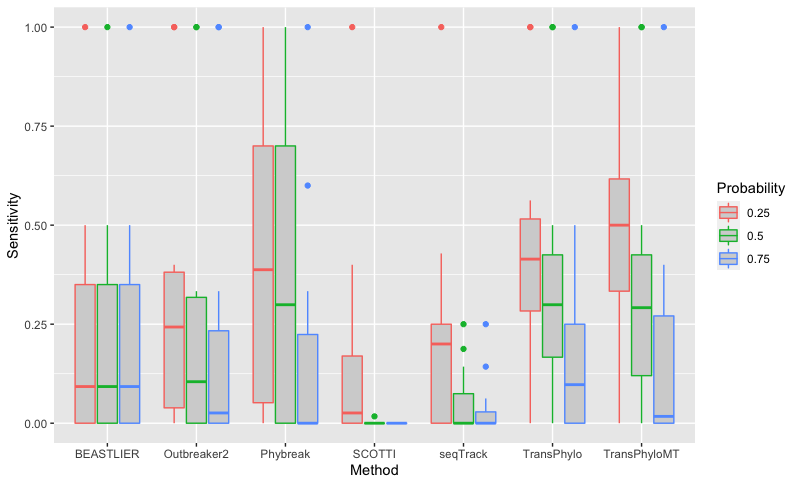

### Figure S2.tiff

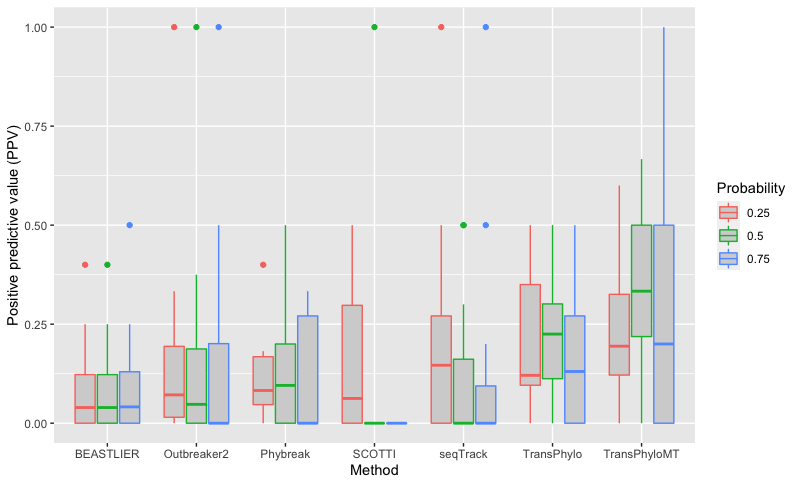

### Figure S3.tiff

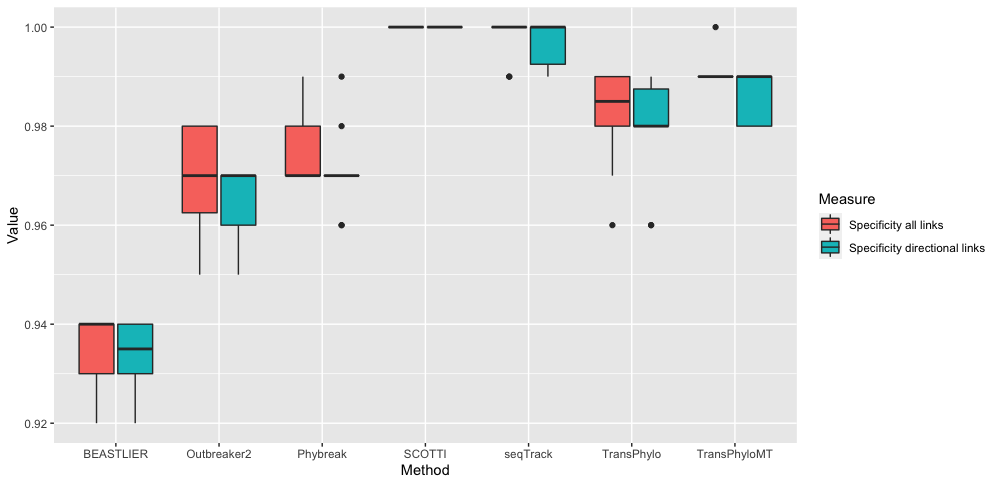
